# Resistance mutation supply modulates the benefit of CRISPR immunity against virulent phages in *Pseudomonas aeruginosa*

**DOI:** 10.1101/2025.10.05.680503

**Authors:** Rosanna CT Wright, Simon C Lovell, Anna Richmond, Christian Harrison, Eleri Ashworth, Aras Kadioglu, Joanne L Fothergill, Ville-Petri Friman, Edze R Westra, Michael A Brockhurst

**Affiliations:** Division of Evolution, Infection and Genomics, School of Biological Sciences, Faculty of Biology, Medicine and Health, University of Manchester, Michael Smith Building, Dover Street, Manchester, M13 9PT, United Kingdom; Environment and Sustainability Institute, Centre for Ecology and Conservation, University of Exeter, Penryn, Cornwall, TR10 9FE, United Kingdom; Department of Clinical Infection, Microbiology and Immunology, University of Liverpool, Liverpool, UK; Department of Microbiology, University of Helsinki, 00014, Helsinki, Finland

## Abstract

Only a fraction of bacterial genomes encode CRISPR-Cas systems but the selective causes of this variation are unexplained. How naturally virulent bacteriophages (phages) select for CRISPR immunity has rarely been tested experimentally. Here, we show against a panel of genetically and functionally diverse virulent phages that CRISPR immunity was not universally beneficial to *Pseudomonas aeruginosa*, and its fitness effect varied strongly between phages in predictable ways. In addition to mechanisms known to alter the effectiveness of CRISPR immunity, such as encoding a matching spacer or a protective nuclear shell, we show that the fitness effect of CRISPR immunity negatively associated with the probability of evolving receptor-based resistance to the phage via spontaneous mutation. Supply of resistance mutations differed strongly between very closely related lipopolysaccharide-binding phages and was associated with variation at the C-terminus of the tail fibre protein altering residues involved in hydrogen bonding and the predicted binding site. Our results show that CRISPR immunity is more beneficial against virulent phages that are harder to evolve resistance to via receptor mutations, suggesting that virulent phage community composition and diversity will be important drivers of the prevalence of CRISPR immunity.

## Introduction

CRISPR-Cas is a prokaryotic adaptive immune system protecting cells from invading mobile genetic elements [1]. CRISPR-Cas systems are encoded by approximately half of sequenced bacterial genomes, yet abundance and system type vary even within clades [2] suggesting that selection for their gain or loss varies widely in nature. Across natural microbiomes, viral abundance positively correlates with the prevalence of CRISPR-Cas systems [3], suggesting that predation by virulent bacteriophages (phages) may select for CRISPR immunity. However, in lab experiments, CRISPR-Cas systems are often ineffective against naturally virulent phage [4–6], in contrast to the highly efficient acquisition of spacers observed against a small number of temperate phages phages when engineered to become obligately virulent [7–10]. This limited range of model systems has nonetheless been useful for understanding the molecular mechanisms of CRISPR immunity, including how partial spacer matches protect infected hosts through priming [11], and the evasion strategies used by phages to negate CRISPR-Cas, such as nucleus formation [6], anti-CRISPRs [12], and DNA modifications [13,14]. However, studies of a wider diversity of phages, including naturally virulent phages, are required to understand the relative importance of these processes in nature, and to determine whether unknown factors exist that control the impact of CRISPR immunity outside the small number of well-studied bacteria-phage interactions. Filling this knowledge gap will also be of likely clinical importance given that virulent phages are a promising alternative to antibiotics for treating multidrug resistant bacterial infections [15], wherein the distribution of CRISPR-Cas systems is likely to be highly variable among strains [16]. Here, it will be vital to be able to predict not only how CRISPR immunity will act against individual virulent phages, but also against combinations of multiple, diverse virulent phages, as often used therapeutically in phage cocktails [17].

To test the fitness effect of encoding CRISPR immunity we compared the efficacy of a panel of 26 diverse naturally virulent phages in all single and pairwise combinations against isogenic *Pseudomonas aeruginosa* PA14 with or without a functional CRISPR-Cas system. PA14 encodes a well-characterised Type-I-F CRISPR-Cas system, and removal of the *csy3* gene is known to disarm CRISPR immunity, allowing direct comparison of phage infection with or without functional CRISPR immunity. The phage panel varied extensively in genome content and sequence identity and targeted a range of cell-surface receptors including lipopolysaccharide (LPS) and Type IV pilus (T4P) (Table 1 and Figure S1) [18]. Importantly, we included phages within the panel that have properties already known to interact with CRISPR immunity, providing benchmarks against which we could compare the other phages: These included PA14P2 which is targeted by a matching spacer in PA14’s CRISPR spacer array, eliciting CRISPR immunity, and phiKZ which encodes a nuclear shell preventing digestion of phage DNA [19], negating CRISPR immunity. Experiments were conducted using a physiologically relevant host airway mimicking medium (synthetic cystic fibrosis medium; SCFM [20]), and the efficacy of virulent phage treatments was quantified as the reduction in bacterial growth in the presence versus absence of phage [21,22]. A total of 351 phage treatments were used, comprising 26 individual virulent phages and all possible pairwise phage combinations (N = 325), and each treatment was replicated 24-fold to increase the probability of observing rare events, such as bacterial escape from phage infection.

**Table 1.**
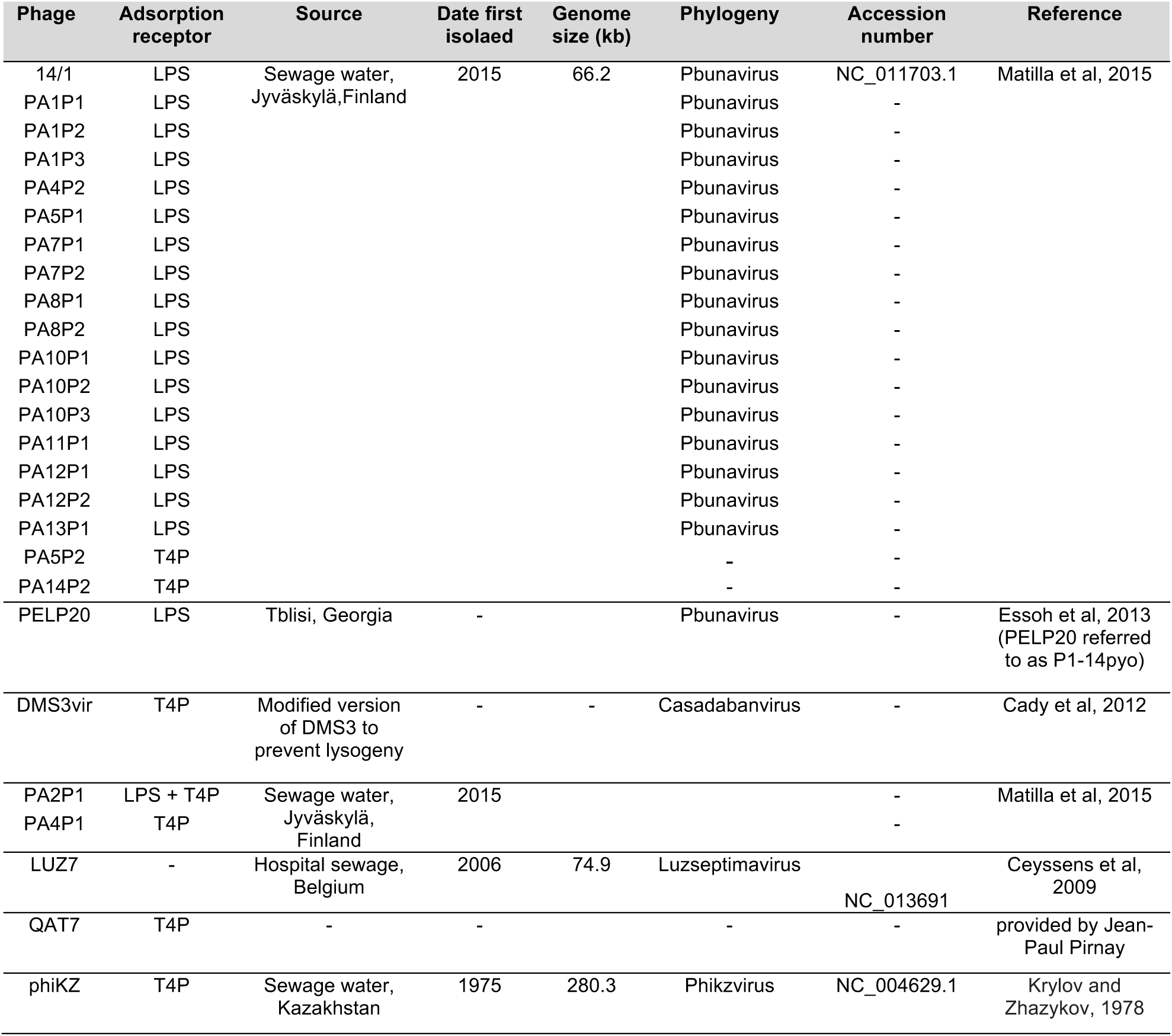
. Phage strain information. . Including known adsorption receptors, isolation source, phylogeny and published genome data.

## Results

### CRISPR immunity is not universally beneficial and varies by phage identity

CRISPR immunity significantly reduced the efficacy of both single and paired phage treatments (Figure 1A, Table S1: CRISPR strain, F_1,16842_=57.03, p<0.00001); efficacy describes the reduction in bacterial growth in the presence versus absence of each phage treatment. Overall, phage pairs had higher efficacy than single phages (Figure 1A; Table S1, number of phages, F_1,16842_=336.22, p<0.00001), and those pairs containing phages both targeting LPS or targeting LPS and T4P achieved highest efficacy (Figure 1C, post-hoc Tukey test on full linear model: all pairwise receptor comparisons p<0.0001 except LPS+T4P vs mixed). However, the efficacy of phage pairs was determined by the inclusion of more effective individual phages rather than synergy between phages, with few phage combinations outperforming their best constituent single phage (Figure 1B). Accordingly, phage identity was the strongest predictor of efficacy overall (Table S1: phage identity, chi-squared=5895, p<0.0001), but individual phages also had a significant interaction with CRISPR immunity (Table S1: PA14P2, 14/1 and PA4P1), suggesting that the host fitness benefit of encoding CRISPR varied against different phages.

**Figure 1.**
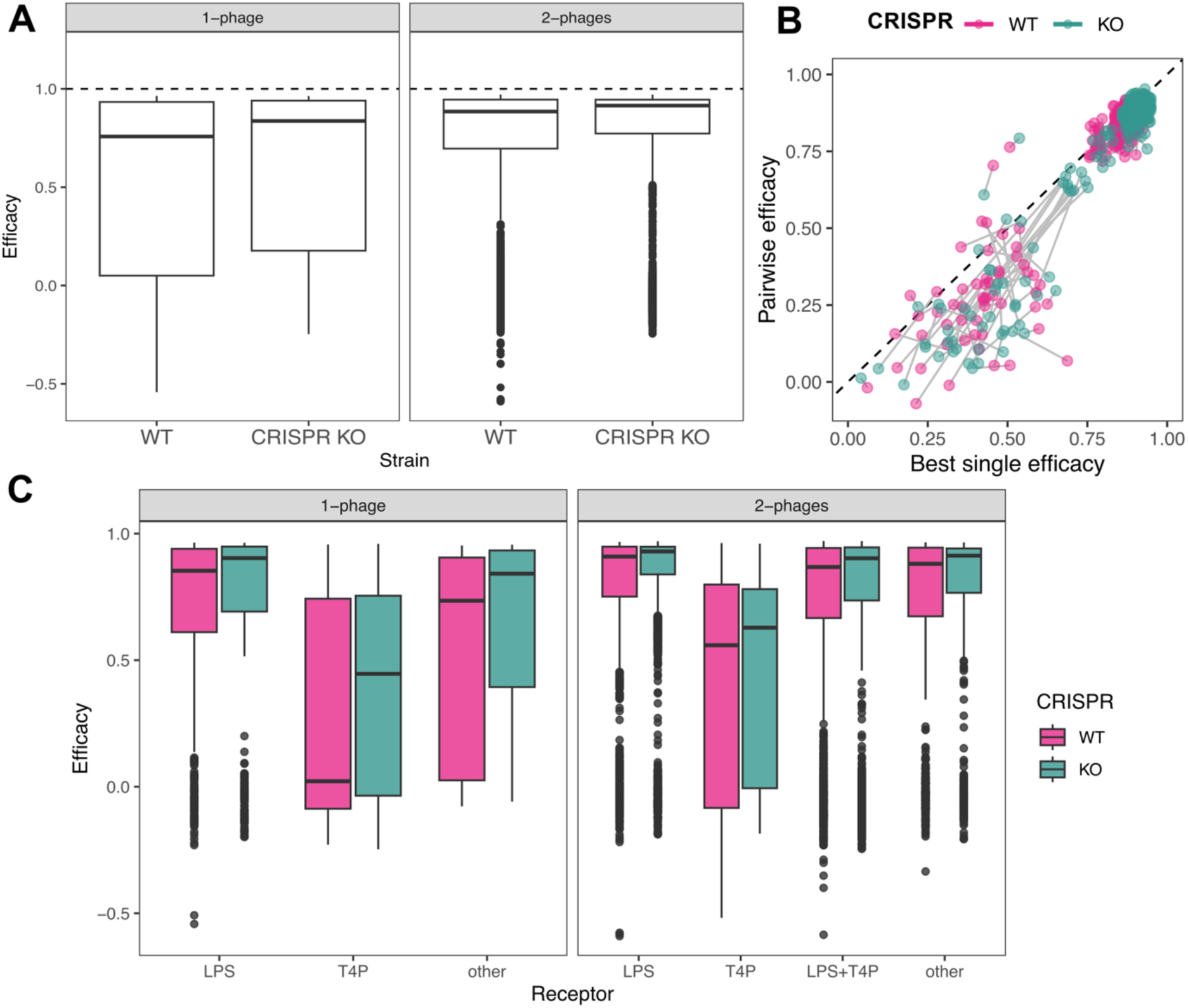
Phage combinations have reduced efficacy in the presence of CRISPR-Cas. **A)** Boxplots showing the difference in efficacy distributions in the presence (WT) and absence (CRISPR KO) of CRISPR immunity in the PA14 background, for single phage treatments (left, N=624 per strain) and 2-phage combinations (right, N=7,800 per strain). **B)** Efficacy of phage pairs in the presence (WT, pink) versus absence (KO, green) of CRISPR immunity is determined by the best single phage present in the pair. Dashed line indicates equal efficacy between best single phage and efficacy of the pair. Data points represent mean of 24 biological replicates; matching phage treatments are joined by a grey line. **C)** Efficacy of phage combinations (left = single phages; right = phage pairs), is influenced by phage receptor target (Receptor) and presence of functional CRISPR-Cas immunity (pink = WT, includes CRISPR-Cas; green = KO, CRISPR-Cas not functional).

To quantify the host fitness benefit of encoding functional CRISPR immunity we next calculated CRISPR effect scores per phage treatment (CRISPR effect = efficacy _CRISPR KO_ – efficacy _WT_). Against individual phages, CRISPR effect scores ranged from positive to negative (Figure 2A). Phage identity, and interaction between phages, were the strongest predictor of CRISPR effect overall (Table S2: phage identity, chi-squared=427.1, p<0.00001; interaction between phages, chi-squared=930.8, p<0.00001). Consistent with their known mechanistic interactions with CRISPR-Cas, the benchmarking phages PA14P2 and phiKZ had the expected positive versus neutral CRISPR effect scores, respectively (Figure 2A, Table S2: PA14P2, F_1,8393_=186.21, p<0.00001 and phiKZ, F_1,8393_=2.49, p=0.115). Observed mean CRISPR effects against individual phages were generally predictive of the CRISPR effects against pairs of these phages (Figure 2B/C; linear model: R^2^=0.227, F_1,323_=96.24, p<0.0001). As such, CRISPR immunity was not universally beneficial against virulent phages or phage combinations but rather benefit depended upon phage identity.

**Figure 2.**
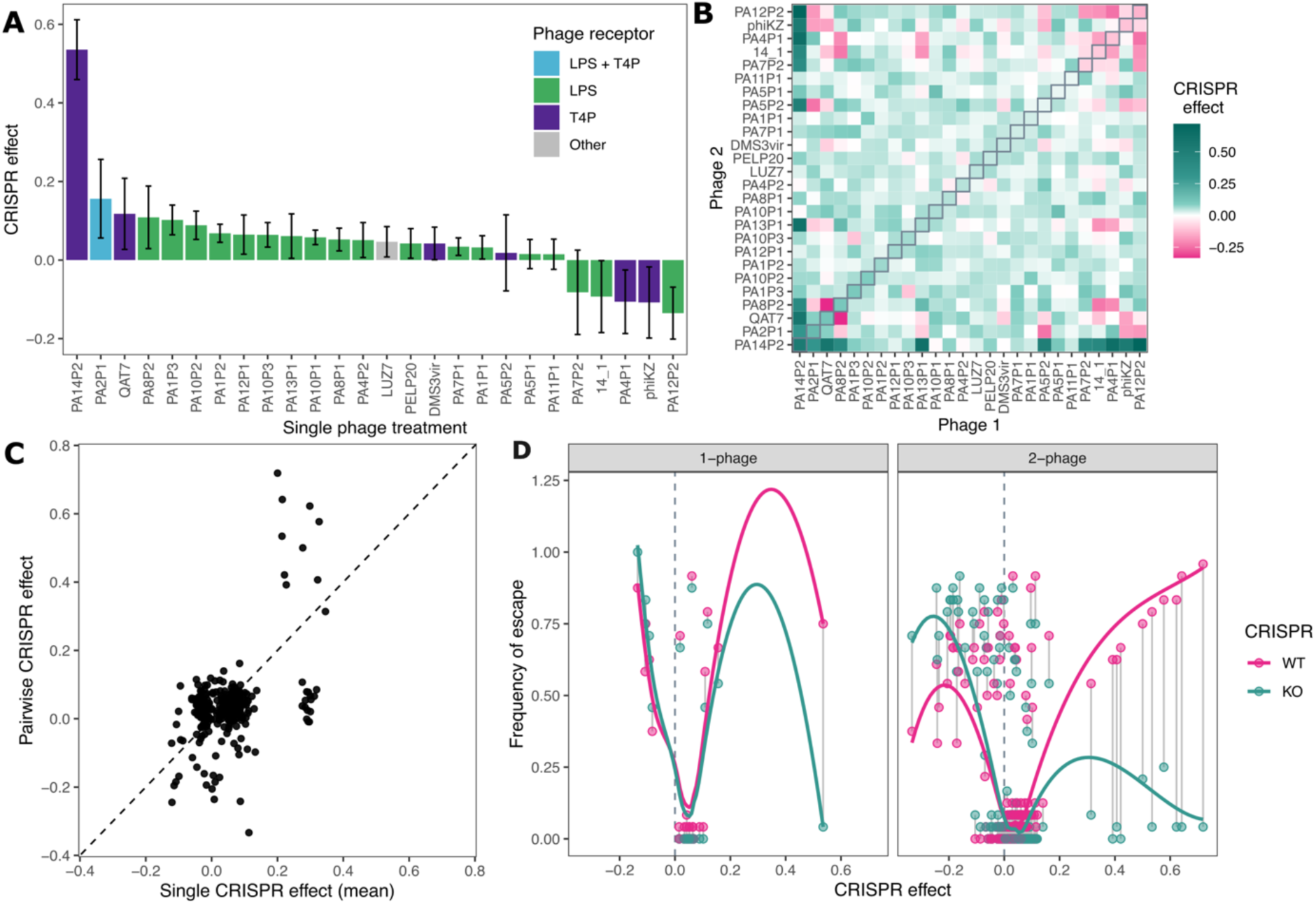
Phage identity determines the impact of CRISPR-Cas immunity on phage efficacy. **A)** Single phage treatments have variable CRISPR effects (efficacy _CRISPR KO_ – efficacy _WT_; mean ± S.E.M, N=24). Bars are ordered by mean CRISPR effect, coloured by phage adsorption receptor (see key). **B**) CRISPR effect against pairs of phages is determined by single-phage CRISPR effects. Heatmap is coloured by mean CRISPR effect of each phage pair (from green to pink representing positive to negative CRISPR effects respectively; N=24). Plots are ordered by the ranking of mean CRISPR effect for individual phages. **C**) Relationship between CRISPR effect of phage pairs and their average individual CRISPR effects. **D**) Relationship between CRISPR effect (mean of 24 replicates) and frequency of bacterial escape, measured as proportion of populations (N=24) which had phage efficacy lower than 0.5 within each treatment. Matching phage treatments are joined by a grey line. Coloured lines show smoothed line of best fit for each bacterial strain.

The observed variation in CRISPR effect between phage treatments could not be explained by known mechanistic interactions with CRISPR-Cas. There was no relationship between CRISPR effect and spacer sequence matches, other than for benchmarking phage PA14P2, nor could we detect any homologs of characterised anti-CRISPR genes in the phage genomes. Furthermore, no acquisition of CRISPR spacers was observed. However, further inspection of individual replicates from the single phage treatments revealed a negative association between CRISPR effect scores and the frequency of bacterial escape from phage infection. Escape was measured as the proportion of replicate populations with <50% reduction in bacterial growth in the presence versus absence of phage (Figure S2). Moreover, both for individual phages and phage pairs, whilst lower CRISPR effect scores resulted in higher frequencies of escape regardless of CRISPR immunity, higher CRISPR effect scores were only associated with high frequencies of escape in the presence of CRISPR immunity (Fig 2D; chi-squared test on interaction between CRISPR effect and CRISPR strain in generalised additive model (GAM) p<0.00001); whilst the highest CRISPR effect scores were associated with the benchmarking phage PA14P2, this interaction between CRISPR effect and CRISPR strain was maintained in the absence of PA14P2 (Figure S3; chi-squared test on GAM interaction p=0.0228). Together these patterns suggest that the benefit of CRISPR immunity declines as the overall probability of bacterial escape increases, but that against phages for which there is a low escape probability, CRISPR immunity enabled increased survival albeit rarely.

### Resistance mutation supply is associated with differences in tail fibre structure

In the absence of a functional CRISPR-Cas system, it is presumed that PA14ΔCRISPR relies upon gain of a surface receptor resistance mutation to escape phage infection. Therefore, we hypothesised that CRISPR immunity becomes more beneficial as the supply of such surface receptor resistance mutations decreases, for example, if fewer viable mutational targets in the LPS biosynthesis pathway alter phage-specific binding sites. To test this, we next quantified resistance mutation supply in the PA14ΔCRISPR background against a subset of 8 closely related PB1-like phages that consisted of 2 groups of phage which varied in CRISPR effect (Kruskal-Wallis: chi-squared=9.32, p=0.0023) and escape frequency (Figure 3A; Kruskal-Wallis: chi-squared=11.36, p-value = 0.0008). Consistent with our hypothesis, both the probability of resistance (Figure 3B; R^2^=0.525, F_1,6_=8.73, p=0.025) and absolute resistance mutation frequencies (Figure 3C; R^2^=0.681, F_1,94_=203.7, p<0.00001) positively correlated with escape frequency in the PA14ΔCRISPR background.

**Figure 3.**
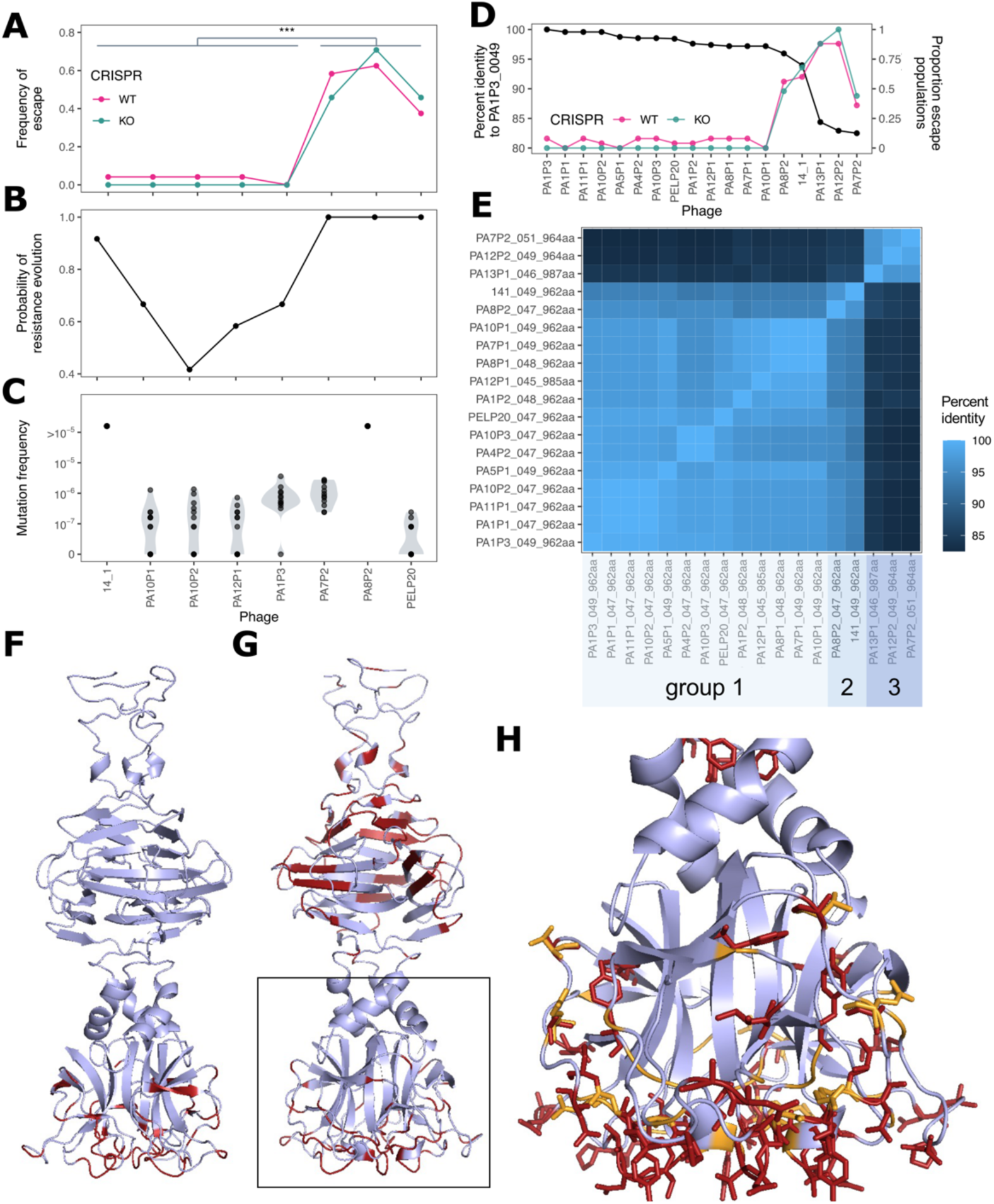
Changes in phage tail structure are associated with reduced availability of *de novo* resistance mutations. **A-C)** For subset of 8 LPS-binding phages: Proportion of populations which ‘escape’ phage infection (N=24) in the presence (WT, pink) versus absence (KO, green) of CRISPR-Cas (A), with statistical significance between phage groups (Kruskal-Wallis test) indicated by stars; Probability of resistance evolution by PA14ΔCRISPR (PA14 *cys3::lacZ*), as proportion of independent replicate populations (N=12) in which resistance evolution was observed (B); Frequency of *de novo* phage resistance mutation by PA14ΔCRISPR (C). **D-E)** For all LPS-binding phages: Proportion of populations which ‘escape’ phage infection (N=24) in the presence (WT, pink) versus absence (KO, green) of CRISPR-Cas relates to changes in a tail protein, gp49 (PA1P3) sequence identitiy (D, black). Equivalent tail proteins were determined via pangenome analysis (*roary*), and percent identity to a representative phage, PA1P3 (gp49), was calculated to provide a measure of divergence in sequence (D/E). Percentage identity across the whole panel of LPS-targeting phages was determined by multiple protein alignment (*clustal omega*; E). Group labels indicate phages sharing the same terminus sequence pattern (Figure S4). **F-H)** Model of the 3D structure of C-terminal portion of phage tail fibre (residues 800-985) for the variants phage 14/1 (F) and phage PA12P2 (G/H) with residues that differ from group 1 highlighted in red and close view (area of G highlighted in grey box shown in H) additionally showing residues that result in loss of hydrogen bond donors or acceptor in orange.

The high nucleotide identity among the PB1-like phages then allowed us to identify possible genetic causes of the variation in resistance mutation supply. Variation in phage tail structures, such as tail fibres, is likely to indicate altered LPS-binding activity and influence the identify of phage-specific surface receptor resistance mutations. We identified a highly variable tail protein whose sequence divergence from a chosen representative (PA1P3’s gp49) varied by up to ∼20% sequence identity across the 18 PB1-like phages (Figure 3D), which explained differences in escape frequency; lower tail protein sequence identity versus PA1P3 gp49 was associated with higher probability of escape in both bacterial strains (Figure 3D). Protein coding sequence alignments for all PB-1-like phages revealed 3 distinct variant groups (Figure 3E & S3): Group 1 resembled PA1P3; Group 2 had amino acid changes between positions 927-978 at the C-terminus of the protein; Group 3 had these changes plus further C-terminus amino acid changes between positions 809-870. Group 1 phages were associated with a low resistance mutation supply, but functional CRISPR immunity allowed rare bacterial escape from phage infection. By contrast, Group 2 and Group 3 phages were associated with higher resistance mutation supply (Figure 3B/C: phages PA8P2, 14/1 and PA7P2), and functional CRISPR immunity had no effect on bacterial escape from phage infection (Figure 3A).

We modelled the tail fibre protein structures for a representative phage from each group: PA1P3 for Group 1, 14/1 for Group 2 and PA12P2 for Group 3. An existing protein structure from a related phage suggests that this tail protein forms a trimer with a beads-on-a-string structure (*20*). All variable residues mapped to the protein surface, with those common to 14/1 and PA12P2 primarily mapping to the C-terminal tip of the tail fibre (Figure 3F-H), which is involved in binding the LPS in *Pbunavirus* phages [23]. These amino acid changes had only very limited effects on charge (only a single charge change event was observed between 14/1 and PA12P2). In contrast, there were large changes to donor/acceptor hydrogen-bonding groups among the phages: 14/1 had 6 fewer donor/acceptor groups than PA1P3 or a total of 18 fewer sites per trimeric tail fibre molecule, whereas PA12P2 had 9 fewer or 27 fewer per trimeric tail fibre molecule (Figure 3H). This suggests changes in the C-terminus of the phage tail fibre may alter the binding target within LPS and hence the genes within the LPS biosynthesis pathway where mutations provide resistance against these phages.

Both 14/1 and PA12P2 are known to bind to the LPS O-antigen, with mutations in the corresponding polymerase, *wzy*, providing resistance to both of these phages [18,24]. To identify the binding receptor of PA1P3, we sequenced 5 independent spontaneous resistant mutants. Four out of five PA1P3-resistant mutants had single missense variants within the L-rhamnose biosynthesis pathway targeting either *rmlA* or *rmlB*, and the final mutant had 2 missense variants targeting *wapR* (rhamnosyltransferase) and *algC* (phosphomannomutase; core LPS biosynthesis) (Table S3). These distinct mutational targets support our hypothesis that PA1P3 binds to a different region of the LPS than 14/1 or PA12P2, most likely a rhamnose residue within the LPS core. To further test this hypothesis, we performed a longer-term evolution experiment wherein replicate populations of PA14ΔCRISPR were propagated by daily serial transfer for 7-days with phages from Group 1, PA1P3 or PELP20, or from Group 2, 14/1 or PA8P2. Resistance evolved against each of the phages (Figure S5) and for one random resistant colony per replicate population we obtained a genome sequence. 10 out of 12 evolved clones had gained mutations in their LPS operons (Figure S6, Table S4). Consistent with phage groups binding to different LPS-regions, these mutations were exclusively in genes encoding the LPS core for clones evolved against Group 1 phages or the O-antigen for clones evolved against Group 2 phages. Specifically, clones resistant to PELP20 gained a mutation in either *rmlA*, encoding a glucose 1-phosphatase thimydyltransferase, or *rmlB*, encoding a dTDP-glucose 4.6-dehydratase, and clones resistant to PA1P3 gained a mutation in *rmlA* or EIP97_RS27925, encoding a putative glycosyltransferase within the same LPS core operon. By contrast, clones resistant to 14/1 or PA8P2 had gained mutations in EIP97_R509780, encoding an O-antigen polymerase. Taken together, our findings suggest that altered hydrogen-bonding at the C-terminus of the phage tail fibre resulted in phage binding to different target sites within the LPS and thus distinct sets of LPS biosynthesis genes wherein mutations provide resistance, resulting in variation in resistance mutation supply.

## Discussion

In this study, we provide experimental evidence for an evolutionary modulator of the fitness benefit of CRISPR immunity. The fitness effect of encoding CRISPR immunity varied by phage identity and was negatively associated with the rate of bacterial escape from phage predation. Thus, CRISPR immunity was more beneficial against virulent phages for which resistance mutation supply was more limited and the probability of evolving resistance was lower, consistent with theoretical predictions [25]. Even in such cases, however, CRISPR-mediated escape was a rare event, occurring in only a fraction of replicate populations. Together these patterns imply that CRISPR-Cas systems are not highly effective against virulent phages in *P. aeruginosa*. Nonetheless, CRISPR immunity may be favoured in communities with higher phage diversity, due to the increased likelihood of encountering virulent phages against which resistance mutations are rare. Our findings parallel ecological studies showing that CRISPR immunity is more beneficial in low resource environments that limit bacterial population size and thus constrain total mutational supply [8,26]. Genetic differences among virulent phages that further limit the supply of resistance mutations would likely be exacerbated in low resource conditions, more strongly favouring CRISPR immunity.

The variation in resistance mutation supply between LPS-binding phages was explained by functional differences in their tail fibre proteins. Specifically, phages that were harder to evolve resistance against (Group 1) had more acceptor/donor hydrogen bonding groups in the C-terminal tip of the tail fibre that interacts with the bacterial cell surface. Protein-polysaccharide interactions are predominantly mediated by hydrogen bonds, suggesting that Group 1 phages may have higher binding affinity for bacterial LPS than those in Group 2 or Group 3. Further, identification of mutational targets within the LPS biosynthesis pathway associated with phage resistance suggest that the molecular binding target for these phages may differ: Group 1 phages were associated with mutations to both L-rhamnose and core LPS biosynthesis, suggesting Group 1 phages may bind to exposed L-rhamnose on the uncapped LPS core, as opposed to the O-antigen for Group 2/3 phages. These results provide greater insight into the mechanisms which drive cross-resistance between phages, whereby asymmetrical cross-resistance between LPS-binding phages [18] may be explained by differences in binding sites across LPS core or O-antigen. Diversity in phage tail structure, even between closely related phages, can thus strongly influence the availability of resistance mutations and the likelihood of broader cross-resistance.

The molecular mechanism by which CRISPR immunity enabled higher frequencies of escape against specific phages remains unclear. No spacer acquisitions could be detected by PCR in the replicates where CRISPR-mediated escape occurred, suggesting a potentially unknown mechanism of protection. Further, it is possible that the csy3 mutation (which removes the functionality of CRISPR-Cas immunity here) could have unforeseen additional effects which impact the comparison between bacterial strains. CRISPR immunity is just one among many antiphage defence systems encoded by bacteria. Indeed, PA14 itself also encodes RM, Gabija and Wadjet [27]; how these and other defences interact together and with CRISPR immunity to provide defence against virulent phage infection is largely unknown [28,29], but such interactions may have affected our results. Moreover, although no anti-CRISPRs were identified in the genomes of phages we used, these proteins are highly variable and poorly characterised, making them difficult to predict by homology-based methods. It is possible therefore that novel ACRs will be discovered in some of these phages in the future, explaining more of the variation in CRISPR effects.

CRISPR-Cas systems are found in ∼40% of *P. aeruginosa* genomes and type IF CRISPR-Cas are notably enriched in isolates from cystic fibrosis lung infections [30], a key clinical target for phage therapy. Whilst we have shown CRISPR immunity is not universally beneficial against single virulent phages, phage therapy often uses cocktails containing multiple, diverse phages. A key benefit of phage cocktails is the potential to limit resistance evolution via mutation, however, our results suggest increasing phage diversity may inadvertently select for pathogens encoding CRISPR immunity. Advances in whole genome sequencing make genetic profiling of patient infections more accessible, to enable identification of bacterial defences likely to hinder phage therapy. However, our data suggest that understanding the availability of resistance mutations during treatment and how this is influenced by phage genetic and environmental factors will be an important additional factor for predicting how CRISPR immunity will impact the efficacy of phage therapy, enabling development of more robust and efficacious phage-based treatment options.

## Acknowledgements

We would like to thank Matti Jalasvuori, Jean-Paul Pirnay and Rob Lavigne for providing phage strains.

## Data Availablility

All experimental data is available in Supporting Information files and raw data is also published on Figshare (doi: 10.6084/m9.figshare.33071714). All sequencing data is published on the European Nucleotide Archive (study accession PRJEB122815).

## Materials and Methods

### Bacteria and phage strains

To understand the impact of CRISPR-Cas, we utilised a well characterised model-system [7] consisting of *Pseudomonas aeruginosa* strain UCBP-PA14 and the isogenic mutant *csy3::LacZ* (herein PA14ΔCRISPR or CRISPR KO), in which the type IF CRISPR-Cas system is not functional. A collection of 26 phage strains were used in this study, chosen due to their ability to infect PA14 and for their genetic and functional diversity (i.e., adsorption receptor usage; Table 1). Phage infection of PA14 was confirmed as ability to produce defined plaques on PA14ΔCRISPR lawn and/or reduce bacterial density of PA14ΔCRISPR in liquid culture (King’s B media) during a 24h growth curve whereby absorbance was measured (OD 600nm) every 20min for 24h at 37°C (Figure S7). Phage adsorption receptors were identified by loss of plaquing ability on bacterial receptor mutants: transposon insertion mutants PAO1::*wzy* (lipopolysaccharide O-antigen biosynthesis) and PAO1::*pilB* (type IV pilus motor protein) were provided by the Washington 2-allelle library [31] and plaquing on these hosts was compared to wild-type PAO1 which was susceptible to all 26 phages.

### Impact of CRISPR-Cas immunity on phage treatment efficacy

To investigate the impact of CRISPR immunity on the efficacy of virulent phages, we challenged the isogenic PA14 strains (with or without a functional CRISPR-Cas system) against 351 different phage treatments. Phage treatments included all single phages (N=26) plus all possible pairwise combinations (N=325). Phages were amplified in host PA14ΔCRISPR, then filter purified (0.22μm filter) and standardised to ∼10^9^PFUml^-1^ (plaque forming units). Phage treatments were prepared in 96-well plate layouts, with pairwise combinations made by mixing equal volumes of standardised phage preparations.

Bacteria were cultured in synthetic cystic fibrosis medium (SCFM; *11*), without the addition of mucin, with individual colonies selected to found 24 individual replicate populations per strain. After overnight growth (37°C, 180rpm), bacterial cultures were sub-cultured into 384-well plates (1 whole plate per replicate strain) to a final density of ∼10^3^ cells per well in fresh SCFM. Phage treatments were added to each plate to give a final multiplicity of infection ∼0.1 (i.e., a ratio of 10 bacteria per phage); each 384-well plate included all phage treatments (N=351) plus negative controls (i.e., no-phage and/or no-bacteria wells). Assay plates were incubated at 37°C, static, with 80% humidity, in sets of 10 384-well plates (replicate batches) for 18h before reading optical density at 600nm to determine bacterial growth.

Phage treatment efficacy was calculated as bacterial growth (OD_600_ at T=18h; Figure S8) in the presence of each phage treatment relative to phage-free growth (Equation 1); average values for phage-free growth were taken within each plate to control for plate and replicate effects. All statistical analysis was performed in R (v4.6.1; [32]). Phage effiacay was tested using a linear mixed effects model (Table S1; using *lmer* function from R package lme4 [33]) testing how efficacy varies by CRISPR strain (i.e., WT vs KO), number of phages (i.e., single or pairs), phage receptor group (LPS-only, T4P-only, other, or mixed), phage identity (as presence/absence matrix for all phages), and interactions between factors. Each effect was tested in a stepwise model, using forward selection and carrying over residuals from each model, to provide in-model test output to explore impact within levels (e.g., individual phages) and compared at model level (chi-squared test) to give overall significance. Replicate batch was included as a random effect. Additional pairwise comparisons within specific factors (e.g., phage receptor) were conducted using a post-hoc Tukey test on the linear model output.

For each bacteria-phage replicate population:

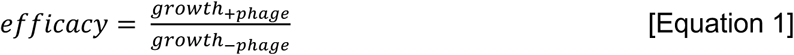

To quantify the impact of CRISPR immunity, the CRISPR Effect (Equation 2) was calculated as the difference between phage efficacy in the absence minus the presence of CRISPR; calculations were performed within replicate, values shown indicate mean ± S.E.M across 24 replicate populations. Significant drivers of the CRISPR effect were assessed using a linear mixed effects model (Table S2; using *lmer* function from R package lme4 [33]) testing how CRISPR effect varies by number of phages (i.e., single or pairs), phage receptor group (LPS-only, T4P-only, other, or mixed), phage identity (as presence/absence matrix for all phages), and interactions between factors. Each effect was tested in a stepwise model, using forward selection and carrying over residuals from each model, to provide in-model test output to explore impact within levels (e.g., individual phages) and compared at model level (chi-squared test) to give overall significance. Replicate batch was included as a random effect. The predictive relationship between CRISPR effect of single phage versus phage pairs was tested using a linear regression.

For each phage treatment (single and pairwise combinations):

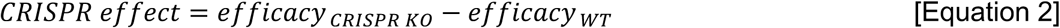

Escape from phage killing was determined at a breakpoint of efficacy equal to 0.5, i.e., less than a 50% reduction in bacterial growth, based on visual inspection of the distribution of efficacy values (Figure S2). The frequency of escape was calculated as the proportion of replicates per treatment which had efficacy lower than 0.5. The relationship between CRISPR effect and frequency of escape was modelled using a generalised additive model (R package *mcgv* [34]), with CRISPR strain fitted both as a main effect and interaction with smoothed CRISPR effect; a chi-squared test was used to interrogate the interaction between CRISPR effect and CRISPR strain.

Colony PCR was used to determine if escape was a result of acquisiton of new spacers within either of PA14’s CRISPR arrays (Table 2) using previously described methods [35]. No acquisition of CRISPR spacers was observed following screening of whole replicate populations and individual colonies selected from each single-phage treatment where escape was observed during the phage efficacy screen.

**Table 2.**
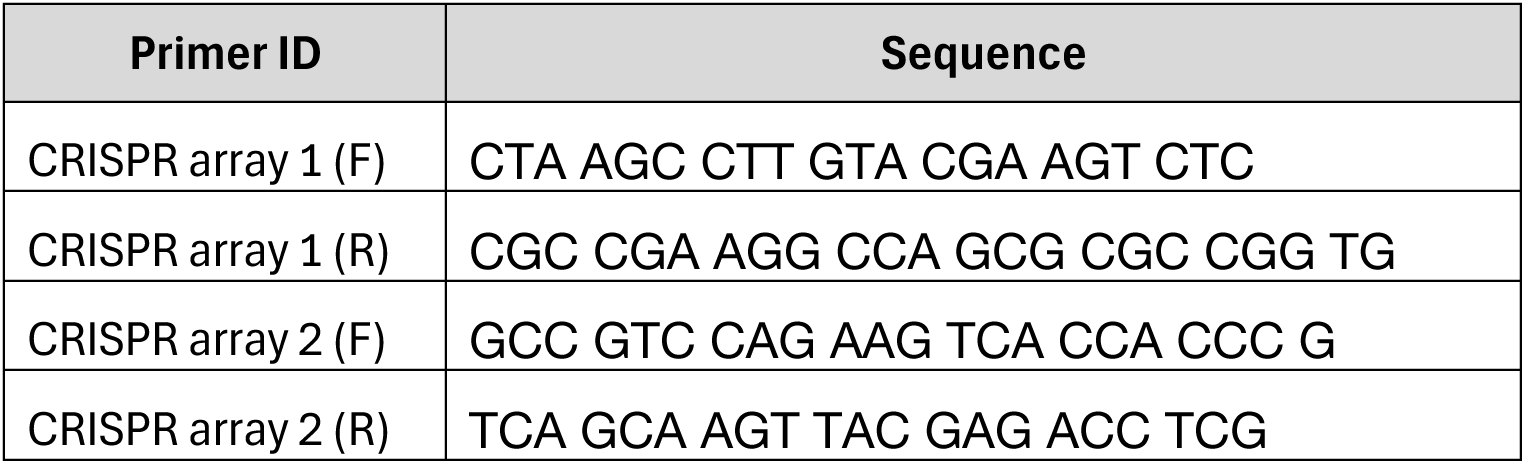
Primer sequence for PA14 CRISPR arrays to detect acquisition of spacers.

### Availability of *de novo* resistance mutations

To calculate the availability of *de novo* resistance mutations, we performed a modified fluctuation test challenging PA14ΔCRISPR against a set of 8 pbunavirus phages which contained 2 distinct patterns of CRISPR effect response and against which varying frequency of escape was observed (statistical significance measured by Kruskal-Wallis test). As CRISPR is not functional in this strain, phage resistance indicates acquired mutations within cell surface receptors preventing phage adsorption. Individual colonies of PA14ΔCRISPR selected to found 12 independent replicates which were then sub-cultured into 8 populations to generate diversity and grown in M9 with 20mM glucose for 18h at 37°C. Independent bacterial subpopulations were then diluted into phage cultures to give a multiplicity of infection of ∼10 phage per bacteria, with 12 independent replicates per phage treatment, mixed and incubated at 37°C for 30min to permit phage adsorption. Mixtures and phage-free controls were then plated and incubated overnight at 37°C before final counts were taken. The probability of resistance evolution was calculated as the proportion of replicates for which any colonies grew after phage challenge. Mutation frequency was calculated as the relative bacterial density (colony forming units per ml) in the presence versus absence of phage. Correlation in probability of resistance evolution and mutation frequency with frequency of escape from phage killing in the PA14ΔCRISPR background was tested with a linear regression

Five independent spontaneous mutants, in PA14ΔCRISPR background, selected for resistance to phage PA1P3 were whole genome sequenced to identify resistance mutations. DNA was extracted from clonal overnight cultures using Wizard Genomic DNA Purification Kit (Promega, Cat. No.: A1120) according to the manufacturer’s instructions, then sequenced by Plasmidsaurus Bacterial Genome Sequencing service using Oxford Nanopore long read technology. Raw reads were aligned to UCBPP-PA14 reference genome (accession NZ_CP034244.1) with minimap2 [36] then filtered for mapping quality and indexed using samtools [37] before variants were called by longshot [38] and annotated using SNPeff [39]. Variants were filtered by quality (read depth >10x coverage; proportion of alternate reads > 0.8), compared to variants identified in control sequence (phage-sensitive PA14 ancestor) to exclude variants which differ from reference genome or common sequencing errors, and verified visually using igv viewer [40].

### Phage whole genome sequencing

For whole genome sequencing, phage DNA was extracted from high-titre phage stocks (∼10^9^ PFUml^-1^) using a phenol-chloroform method described previously [41]. Library preparation followed by paired end short-read illumina sequencing on MiSeq was performed by Centre for Genomic Research, University of Liverpool. Phage genomes were assembled using spades [42], then annotated using pharokka [43] using default parameters and location of the large terminase subunit to reorient phage genomes for comparison. A pangenome of the 18 Pbunavirus phages (Table 1) was constructed using roary [44]. Phage genomes were searched for the presence of anti-CRISPRs using AcaFinder [45], but no anti-CRISPR genes were identified except for the previously described [46] anti-CRISPR genes present in DMS3*vir* which are not active against PA14’s IF CRISPR system. Presence of CRISPR spacers in PA14 matching our phage collection were identified using CRISPR Target [47]: only spacers matching PA14P2 were identified, corresponding to perfect matches with 3 spacer sequences.

### Protein Modelling

Comparison of the phage tail proteins homologous to PA1P3 gp49 across the collection of 18 Pbunavirus genomes was made using clustal omega [48] to align coding sequences, then visualised using Jalview [49]. Protein structures were modelled with the alphafold3 server [50] using default parameters. Structures were modelled as homo-trimers. Since all of the sequence variation is in the region of residues 800-985 only this region was modelled. Types of amino-acid change were identified by inspection.

### Resistance evolution assay

Overnight cultures of PA14ΔCRISPR were diluted 1:100 into fresh KB medium and infected with one of four phages (14-1, PA8P2, PELP20, PA1P3) at an initial MOI of 0.01, with 3 biological replicates per phage. Infected cultures were incubated at 37°C with shaking at 180 rpm and 1% diluted into fresh medium every 24 hours for 7 days. After 7 transfers, cultures were plated out for single colonies and 12 colonies from each treatment were picked to test for phage resistance.

Overnight cultures of putative resistant colonies were diluted 1:100 into a 96-well plate both with and without the phage against which they were evolved at an MOI of 100 with colonies of the ancestral strain included as control. Plates were incubated with shaking at 37°C for 24 hours with OD600 measured every 10 minutes. The area under the curve (AUC) was calculated for each sample and the ratio of AUC (phage treated) / AUC (phage free) used to determine phage resistance with a score >0.8 indicated phage resistance. A single phage resistant clone from each population was selected at random for sequencing with Plasmidsaurus whole genome sequencing to screen for mutations.

Raw reads were mapped to the UCBPP-PA14 reference genome (accession NZ_CP034244.1) with minimap2 [36]07/10/2026 14:52:00, filtered and indexed with samtools [37], variants called with Clair3 using model r1040_e82_400bps_sup_v430_bacteria_finetuned [51] and annotated with SNPeff [35]. Background mutations common to all samples were removed with bcftools [52] and variants of low quality were removed manually (including any mutations in *csy3*, known to be mutated in the reference genome by *lacZ* insertion).

