## Supplementary Information for "Resistance mutation supply modulates the benefit of CRISPR immunity against virulent phages in *Pseudomonas aeruginosa*"

| | | | Chi-squared Test | | | In-model test | | | |
|---|---|---|---|---|---|---|---|---|---|
| **Model** | **Model Equation** | **AIC** | **DF** | **ChiSq** | **p-value** | **Test** | **DF** | **F-value** | **p-value** |
| **0** | Efficacy ~ (1\|Batch) | 8768 | | | | | | | |
| **1** | residuals(model 0) ~ CRISPR Strain + (1\|Batch) | 8692 | 1 | 78.3 | **<0.00001** | | 16842 | 57.03 | **<0.00001** |
| **2** | residuals(model 1) ~ number of phages + (1\|Batch) | 8359 | 0 | 332.9 | | | 16842 | 336.22 | **<0.00001** |
| **3** | residuals(model 2) ~ phage receptor group (i.e., LPS, T4P or combined) + (1\|Batch) | 6839 | 3 | 1525.9 | **<0.00001** | | 16839 | 399.17 | **<0.00001** |
| **4** | residuals(model 3) ~ interaction between number of phages and phage receptor group + (1\|Batch) | 6796 | 3 | 48.7 | **<0.00001** | | 16836 | 6.96 | **<0.00001** |
| **5** | residuals(model 4) ~ phage identity (i.e., presence/absence) + (1\|Batch) | 940 | 19 | 5894.7 | **<0.00001** | QAT7 | 16817 | 48.39 | **<0.00001** |
| | | | | | | LUZ7 | 16817 | 146.43 | **<0.00001** |
| | | | | | | PA5P2 | 16817 | 15.70 | **0.00007** |
| | | | | | | phiKZ | 16817 | 9.67 | **0.00187** |
| | | | | | | PA1P1 | 16817 | 84.39 | **<0.00001** |
| | | | | | | DMS3vir | 16817 | 358.96 | **<0.00001** |
| | | | | | | PELP20 | 16817 | 65.94 | **<0.00001** |
| | | | | | | PA1P2 | 16817 | 79.34 | **<0.00001** |
| | | | | | | PA1P3 | 16817 | 21.06 | **<0.00001** |
| | | | | | | PA2P1 | 16817 | 136.40 | **<0.00001** |
| | | | | | | PA4P1 | 16817 | 78.18 | **<0.00001** |
| | | | | | | PA4P2 | 16817 | 89.72 | **<0.00001** |
| | | | | | | PA5P1 | 16817 | 51.18 | **<0.00001** |
| | | | | | | PA7P1 | 16817 | 37.26 | **<0.00001** |
| | | | | | | PA7P2 | 16817 | 346.74 | **<0.00001** |
| | | | | | | PA8P1 | 16817 | 50.78 | **<0.00001** |
| | | | | | | PA8P2 | 16817 | 163.93 | **<0.00001** |
| | | | | | | PA10P1 | 16817 | 25.80 | **<0.00001** |
| | | | | | | PA10P2 | 16817 | 74.97 | **<0.00001** |
| | | | | | | PA10P3 | 16817 | 74.73 | **<0.00001** |
| | | | | | | PA11P1 | 16817 | 68.03 | **<0.00001** |
| | | | | | | PA12P1 | 16817 | 69.34 | **<0.00001** |
| | | | | | | PA12P2 | 16817 | 565.91 | **<0.00001** |
| | | | | | | PA13P1 | 16817 | 613.54 | **<0.00001** |
| | | | | | | PA14P2 | 16817 | 0.23 | 0.63149 |
| | | | | | | 14/1 | 16817 | 247.33 | **<0.00001** |
| **6** | residuals(model 5) ~ Interaction between CRISPR strain + phage identity + (1\|Batch) | 727 | 1 | 0.0 | 1 | Strain:QAT7 | 16791 | 0.88 | 0.41338 |
| | | | | | | Strain:LUZ7 | 16791 | 0.04 | 0.96008 |
| | | | | | | Strain:PA5P2 | 16791 | 0.37 | 0.68874 |
| | | | | | | Strain:phiKZ | 16791 | 0.70 | 0.49798 |
| | | | | | | Strain:PA1P1 | 16791 | 0.19 | 0.82331 |
| | | | | | | Strain:DMS3vir | 16791 | 1.38 | 0.25105 |
| | | | | | | Strain:PELP20 | 16791 | 0.02 | 0.97556 |
| | | | | | | Strain:PA1P2 | 16791 | 0.01 | 0.99327 |
| | | | | | | Strain:PA1P3 | 16791 | 0.27 | 0.76115 |
| | | | | | | Strain:PA2P1 | 16791 | 0.63 | 0.53016 |
| | | | | | | Strain:PA4P1 | 16791 | 3.03 | **0.04852** |
| | | | | | | Strain:PA4P2 | 16791 | 0.56 | 0.56904 |
| | | | | | | Strain:PA5P1 | 16791 | 0.00 | 0.99869 |
| | | | | | | Strain:PA7P1 | 16791 | 0.19 | 0.82345 |
| | | | | | | Strain:PA7P2 | 16791 | 0.14 | 0.86863 |
| | | | | | | Strain:PA8P1 | 16791 | 0.02 | 0.98176 |
| | | | | | | Strain:PA8P2 | 16791 | 0.42 | 0.65828 |
| | | | | | | Strain:PA10P1 | 16791 | 0.48 | 0.62059 |
| | | | | | | Strain:PA10P2 | 16791 | 0.04 | 0.95645 |
| | | | | | | Strain:PA10P3 | 16791 | 0.35 | 0.70270 |
| | | | | | | Strain:PA11P1 | 16791 | 0.22 | 0.80070 |
| | | | | | | Strain:PA12P1 | 16791 | 0.11 | 0.89878 |
| | | | | | | Strain:PA12P2 | 16791 | 1.05 | 0.35092 |
| | | | | | | Strain:PA13P1 | 16791 | 0.08 | 0.91876 |
| | | | | | | Strain:PA14P2 | 16791 | 117.64 | **<0.00001** |
| | | | | | | Strain:14/1 | 16791 | 6.96 | **0.00095** |
| **7** | residuals(model 6) ~ Interaction between number of phages + phage identity + (1\|Batch) | 100 | 25 | 889.5 | **<0.00001** | Number:QAT7 | 16792 | 6.03 | **0.00242** |
| | | | | | | Number:LUZ7 | 16792 | 7.31 | **0.00067** |
| | | | | | | Number:PA5P2 | 16792 | 2.78 | 0.06201 |
| | | | | | | Number:phiKZ | 16792 | 0.91 | 0.40301 |
| | | | | | | Number:PA1P1 | 16792 | 5.60 | **0.00369** |
| | | | | | | Number:DMS3vir | 16792 | 20.31 | **<0.00001** |
| | | | | | | Number:PELP20 | 16792 | 3.69 | **0.02507** |
| | | | | | | Number:PA1P2 | 16792 | 4.68 | **0.00929** |
| | | | | | | Number:PA1P3 | 16792 | 3.52 | **0.02975** |
| | | | | | | Number:PA2P1 | 16792 | 5.96 | **0.00258** |
| | | | | | | Number:PA4P1 | 16792 | 5.99 | **0.00251** |
| | | | | | | Number:PA4P2 | 16792 | 5.18 | **0.00564** |
| | | | | | | Number:PA5P1 | 16792 | 5.05 | **0.00641** |
| | | | | | | Number:PA7P1 | 16792 | 3.99 | **0.01844** |
| | | | | | | Number:PA7P2 | 16792 | 3.81 | **0.02212** |
| | | | | | | Number:PA8P1 | 16792 | 4.77 | **0.00848** |
| | | | | | | Number:PA8P2 | 16792 | 12.92 | **<0.00001** |
| | | | | | | Number:PA10P1 | 16792 | 6.50 | **0.00151** |
| | | | | | | Number:PA10P2 | 16792 | 4.82 | **0.00807** |
| | | | | | | Number:PA10P3 | 16792 | 4.15 | **0.01570** |
| | | | | | | Number:PA11P1 | 16792 | 4.70 | **0.00913** |
| | | | | | | Number:PA12P1 | 16792 | 3.65 | **0.02601** |
| | | | | | | Number:PA12P2 | 16792 | 90.95 | **<0.00001** |
| | | | | | | Number:PA13P1 | 16792 | 80.66 | **<0.00001** |
| | | | | | | Number:PA14P2 | 16792 | 2.68 | 0.06828 |
| | | | | | | Number:14/1 | 16792 | 46.61 | **<0.00001** |
| **8** | residuals(model 7) ~ Interaction between phages + (1\|Batch) | -5593 | 298 | 6916.0 | **<0.00001** | | | | |

***Table S1. Statistical table for phage efficacy models.*** *Linear mixed effects model output testing how efficacy varies by CRISPR strain (i.e., WT vs KO), Number of phages (i.e., single or pairs), Phage Receptor Group (LPS-only, T4P-only, other, or mixed), Phage identity (as presence/absence matrix for all phages), and interactions between factors. Models were built up stepwise (carrying over residuals) to provide in-model test output to explore impact within levels (e.g., individual phages) and compared at model level (chi-squared test) to give overall significance. Replicate batch was included as a random effect in each model. Non-significant interactions were dropped from the model.*

| | | Chi-squared test | | | | In-model test | | | |
|---|---|---|---|---|---|---|---|---|---|
| **Model** | **Model Equation** | **AIC** | **ChiSq** | **DF** | **p-value** | **Test** | **DF** | **F-value** | **p-value** |
| ce0 | CRISPR Effect ~ 0 + (1\|Batch) | 2018 | | | | | | | |
| ce1 | residuals(model ce0) ~ number of phages +(1\|Batch) | 1996 | 25.37 | 2 | <0.00001 | | | | |
| ce2 | residuals(model ce1) ~ phage receptor group +(1\|Batch) | 1968 | 33.94 | 3 | <0.00001 | | | | |
| ce3 | residuals(model ce2) ~ phage identity +(1\|Batch) | 1585 | 427.13 | 22 | <0.00001 | QAT7 | 8393 | 2.94 | 0.0866 |
| | | | | | | LUZ7 | 8393 | 1.60 | 0.2056 |
| | | | | | | PA5P2 | 8393 | 1.52 | 0.2178 |
| | | | | | | phiKZ | 8393 | 2.49 | 0.1148 |
| | | | | | | PA1P1 | 8393 | 0.18 | 0.6710 |
| | | | | | | DMS3vir | 8393 | 4.08 | **0.0433** |
| | | | | | | PELP20 | 8393 | 1.45 | 0.2286 |
| | | | | | | PA1P2 | 8393 | 0.80 | 0.3724 |
| | | | | | | PA1P3 | 8393 | 2.82 | 0.0931 |
| | | | | | | PA2P1 | 8393 | 0.22 | 0.6354 |
| | | | | | | PA4P1 | 8393 | 7.30 | **0.0069** |
| | | | | | | PA4P2 | 8393 | 0.00 | 0.9949 |
| | | | | | | PA5P1 | 8393 | 1.09 | 0.2955 |
| | | | | | | PA7P1 | 8393 | 2.47 | 0.1157 |
| | | | | | | PA7P2 | 8393 | 0.26 | 0.6098 |
| | | | | | | PA8P1 | 8393 | 1.40 | 0.2363 |
| | | | | | | PA8P2 | 8393 | 0.02 | 0.8753 |
| | | | | | | PA10P1 | 8393 | 3.61 | 0.0576 |
| | | | | | | PA10P2 | 8393 | 1.62 | 0.2027 |
| | | | | | | PA10P3 | 8393 | 0.05 | 0.8216 |
| | | | | | | PA11P1 | 8393 | 0.15 | 0.7002 |
| | | | | | | PA12P1 | 8393 | 2.03 | 0.1543 |
| | | | | | | PA12P2 | 8393 | 0.08 | 0.7790 |
| | | | | | | PA13P1 | 8393 | 0.38 | 0.5353 |
| | | | | | | PA14P2 | 8393 | 186.21 | **<0.00001** |
| | | | | | | 14/1 | 8393 | 5.76 | **0.0164** |
| ce4 | residuals(model ce3) ~ interaction betweeen number of phages and phage identity +(1\|Batch) | 1567 | 67.82 | 25 | <0.00001 | Number:QAT7 | 8368 | 0.48 | 0.6161 |
| | | | | | | Number:LUZ7 | 8368 | 0.02 | 0.9779 |
| | | | | | | Number:PA5P2 | 8368 | 0.56 | 0.5704 |
| | | | | | | Number:phiKZ | 8368 | 5.49 | **0.0041** |
| | | | | | | Number:PA1P1 | 8368 | 0.02 | 0.9756 |
| | | | | | | Number:DMS3vir | 8368 | 0.09 | 0.9153 |
| | | | | | | Number:PELP20 | 8368 | 0.02 | 0.9773 |
| | | | | | | Number:PA1P2 | 8368 | 0.21 | 0.8124 |
| | | | | | | Number:PA1P3 | 8368 | 0.54 | 0.5804 |
| | | | | | | Number:PA2P1 | 8368 | 2.34 | 0.0967 |
| | | | | | | Number:PA4P1 | 8368 | 4.36 | **0.0127** |
| | | | | | | Number:PA4P2 | 8368 | 0.17 | 0.8447 |
| | | | | | | Number:PA5P1 | 8368 | 0.11 | 0.8937 |
| | | | | | | Number:PA7P1 | 8368 | 0.05 | 0.9538 |
| | | | | | | Number:PA7P2 | 8368 | 2.22 | 0.1089 |
| | | | | | | Number:PA8P1 | 8368 | 0.05 | 0.9521 |
| | | | | | | Number:PA8P2 | 8368 | 1.27 | 0.2795 |
| | | | | | | Number:PA10P1 | 8368 | 0.03 | 0.9701 |
| | | | | | | Number:PA10P2 | 8368 | 0.41 | 0.6622 |
| | | | | | | Number:PA10P3 | 8368 | 0.29 | 0.7514 |
| | | | | | | Number:PA11P1 | 8368 | 0.05 | 0.9495 |
| | | | | | | Number:PA12P1 | 8368 | 0.10 | 0.9050 |
| | | | | | | Number:PA12P2 | 8368 | 4.12 | **0.0163** |
| | | | | | | Number:PA13P1 | 8368 | 0.18 | 0.8382 |
| | | | | | | Number:PA14P2 | 8368 | 9.62 | **0.00007** |
| | | | | | | Number:X14_1 | 8368 | 2.03 | 0.1546 |
| ce5 | residuals(model ce4) ~ Interaction between phages +(1\|Batch) | 1235 | 930.80 | 299 | <0.00001 | | | | |

***Table S2. Statistical table for CRISPR effect models.*** *Linear mixed effects model output testing how CRISPR effect varies by Number of phages (i.e., single or pairs), Phage Receptor Group (LPS-only, T4P-only, other, or mixed), Phage identity (as presence/absence matrix for all phages), and interactions between factors. Models were built up stepwise (carrying over residuals) to provide in-model test output to explore impact within levels (e.g., individual phages) and compared at model level (chi-squared test) to give overall significance. Replicate batch was included as a random effect in each model. Non-significant interactions were dropped from the model.*

| Clone | Gene Name | Gene ID | Gene Product | Genome position (bp) | Reference | Alternate | Change | EffectType | Impact | Alternate read proportion | Coverage |
|---|---|---|---|---|---|---|---|---|---|---|---|
| 1_PA1P3 | rmlA (rfbA) | EIP97_RS28795 | glucose-1-phosphate thymidylyltransferase | 6083406 | A | C | His17Pro | missense variant | moderate | 0.94 | 88 |
| 2_PA1P3 | wapR | EIP97_RS27920 | alpha-1,3-rhamnosyltransferase | 5888477 | A | C | Ser97Ala | missense variant | moderate | 0.91 | 102 |
| 2_PA1P3 | algC | EIP97_RS29620 | phosphomannomutase | 6265538 | G | T | Arg836Leu | missense variant | moderate | 0.95 | 87 |
| 3_PA1P3 | rmlB (rfbB) | EIP97_RS28785 | dTDP-glucose 4,6-dehydratase | 6082051 | G | A | Gly219Ser | missense variant | moderate | 0.99 | 73 |
| 4_PA1P3 | rmlA (rfbA) | EIP97_RS28795 | glucose-1-phosphate thymidylyltransferase | 6083522 | A | C | Thr56Pro | missense variant | moderate | 0.94 | 77 |
| 5_PA1P3 | rmlA (rfbA) | EIP97_RS28795 | glucose-1-phosphate thymidylyltransferase | 6083688 | A | C | Asn111Thr | missense variant | moderate | 1.00 | 100 |

***Table S3. Genome variants associated with resistance to phage PA1P3 (spontaneous mutants)***

| Clone | Gene Name | Gene ID | Gene Product | Genome Position | Reference | Alternate | Change | Effect Type | Impact | Alternate Read Percentage | Coverage | Role |
|---|---|---|---|---|---|---|---|---|---|---|---|---|
| 141_1 | rpoB | EIP97_RS03660 | DNA-directed RNA polymerase subunit beta | 746979 | ACCAGCTGGAAAAGGCTGGCGTGAG | A | Lys284-Glu291 (del) | inframe deletion | MODERATE | 96.15 | 130 | Other |
| 141_2 | rpoB | EIP97_RS03660 | DNA-directed RNA polymerase subunit beta | 746979 | ACCAGCTGGAAAAGGCTGGCGTGAG | A | Lys284-Glu291 (del) | inframe deletion | MODERATE | 96.89 | 161 | Other |
| 141_2 | EIP97_RS28520 | EIP97_RS28520 | DUF2339 domain-containing protein | 6029418 | C | T | Gly1043Arg | missense variant | MODERATE | 98.82 | 169 | Other |
| 141_3 | EIP97_RS09780 | EIP97_RS09780 | O-antigen polymerase | 2031710 | G | GT | Tyr46 (fs) | frameshift variant | HIGH | 94.44 | 18 | O-antigen |
| 141_3 | pepN | EIP97_RS10145 | Aminopeptidase N | 2106473 | A | T | Gln514Leu | missense variant | MODERATE | 99.17 | 121 | Other |
| 141_3 | pilM | EIP97_RS28155 | Type IV pilus assembly protein | 5950873 | A | C | Val60Gly | missense variant | MODERATE | 95.59 | 136 | Other |
| PA8P2_1 | EIP97_RS09780 | EIP97_RS09780 | O-antigen polymerase | 2032685 | A | AT | Ala371 (fs) | frameshift variant | HIGH | 95.45 | 22 | O-antigen |
| PA8P2_1 | pilM | EIP97_RS28155 | Type IV pilus assembly protein | 5950873 | A | C | Val60Gly | missense variant | MODERATE | 93.33 | 120 | Other |
| PA8P2_2 | EIP97_RS09780 | EIP97_RS09780 | O-antigen polymerase | 2031710 | GT | G | Tyr46 (fs) | frameshift variant | HIGH | 89.47 | 19 | O-antigen |
| PA8P2_3 | EIP97_RS09780 | EIP97_RS09780 | O-antigen polymerase | 2031710 | G | GT | Tyr46 (fs) | frameshift variant | HIGH | 100 | 10 | O-antigen |
| PELP20_1 | rmlA (rfbA) | EIP97_RS28795 | Glucose-1-phosphate thymidylyltransferase | 6083606 | T | C | Ser84Pro | missense variant | MODERATE | 100 | 25 | Core |
| PELP20_2 | rmlB (rfbB) | EIP97_RS28785 | dTDP-glucose 4,6-dehydratase | 6081631 | A | T | Asn79Tyr | missense variant | MODERATE | 96.61 | 59 | Core |
| PELP20_3 | rmlB (rfbB) | EIP97_RS28785 | dTDP-glucose 4,6-dehydratase | 6082042 | T | TACGGCG | Asp218-Gly219 (ins) | inframe insertion | MODERATE | 69.7 | 165 | Core |
| PA1P3_1 | rmlA (rfbA) | EIP97_RS28795 | Glucose-1-phosphate thymidylyltransferase | 6083606 | T | C | Ser84Pro | missense variant | MODERATE | 98.78 | 245 | Core |
| PA1P3_2 | EIP97_RS27925 | EIP97_RS27925 | Glycosyltransferase family protein | 5889771 | G | A | Gln38* | stop gained | HIGH | 93.44 | 61 | Core |
| PA1P3_3 | lasR | EIP97_RS19295 | Transcriptional regulator | 4085119 | T | G | Asp73Ala | missense variant | MODERATE | 94.92 | 59 | Other |
| PA1P3_3 | rmlA (rfbA) | EIP97_RS28795 | Glucose-1-phosphate thymidylyltransferase | 6083522 | A | C | Thr56Pro | missense variant | MODERATE | 96.47 | 85 | Core |

***Table S4. Evolved genome variants associated with resistance to phages from tail fibre group 1 (PELP20 and PA1P3) and tail fibre group 2 (14/1, PA8P2) during the evolution experiment.***

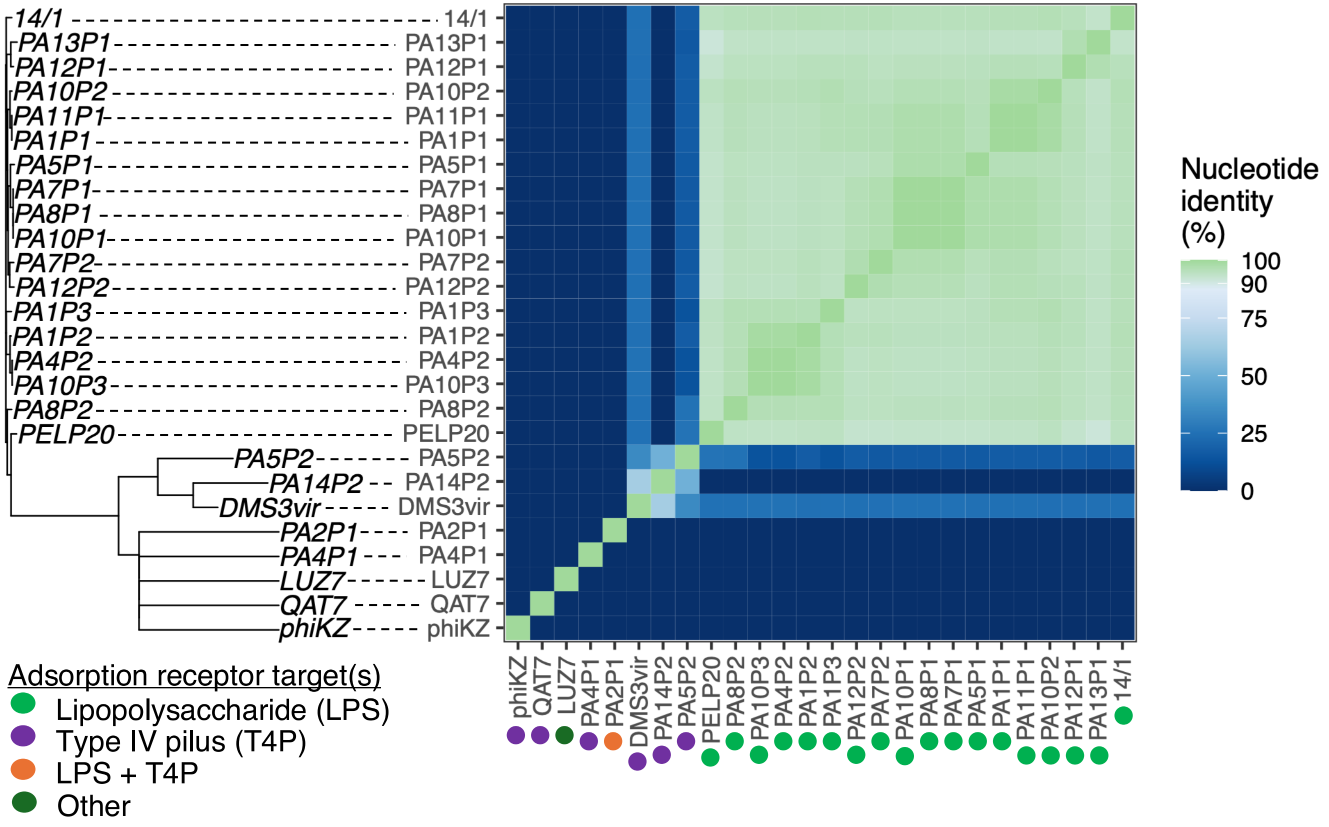


***Figure S1. Phage genetic relatedness.*** *Neighbour joining tree describes phage relatedness based on overall nucleotide identity (% similarity; see heatmap key). Coloured dots indicate phage adsorption receptors (see key). The data underlying this figure can be found in S7 Data.*

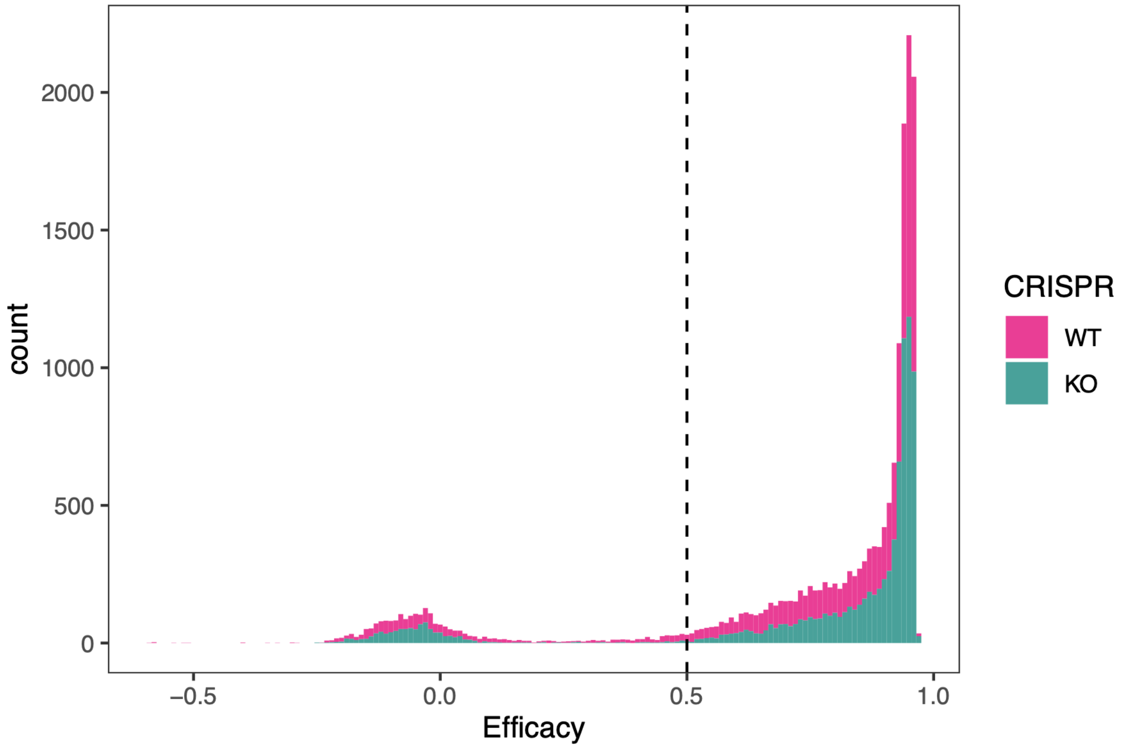


***Figure S2. Distribution of efficacy values to determine breakpoint for escape.*** *Dashed line indicates a breakpoint of efficacy=0.5; populations with efficacy<0.5 were classified as 'escape populations', indicating less than 50% reduction in bacterial growth. The data underlying this figure can be found in S1 Data.*

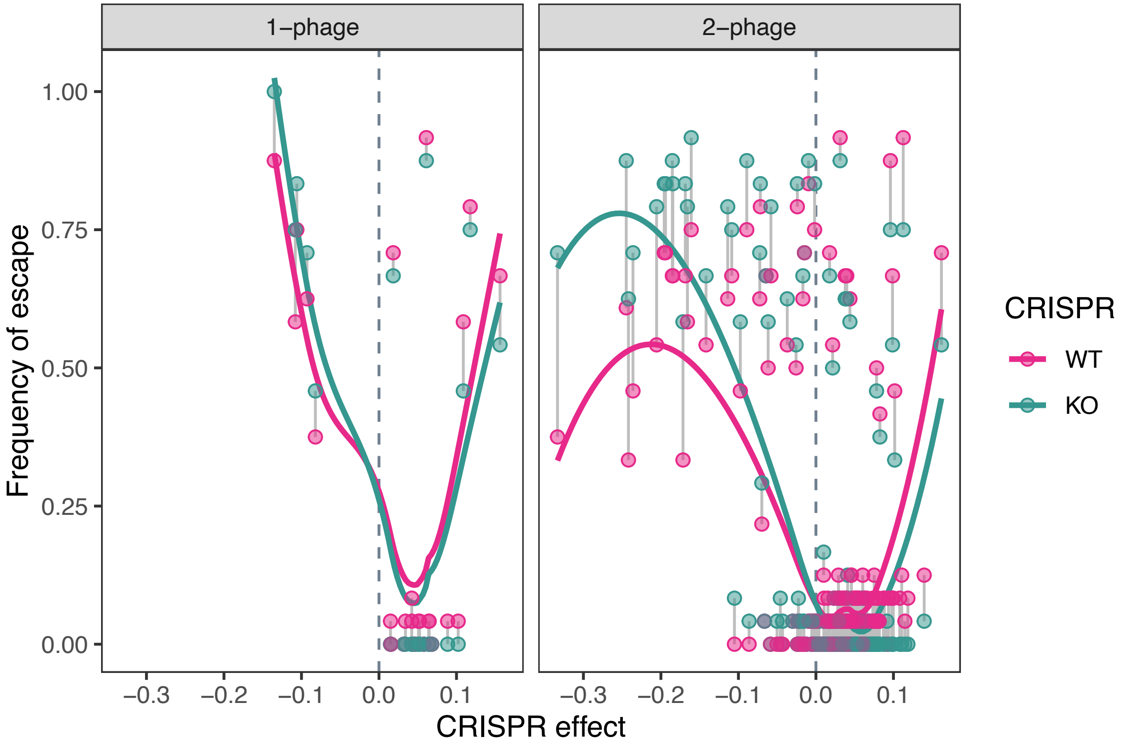


***Figure S3. Relationship between CRISPR effect and Frequency of escape excluding phage treatments containing phage PA14P2.*** *Plotted values for CRISPR effect show mean of 24 replicates; frequency of bacterial escape was measured as proportion of populations which had phage efficacy lower than 0.5 within each treatment. Matching phage treatments are joined by a grey line. Coloured lines show smoothed line of best fit for each bacterial strain. The data underlying this figure can be found in S2 Data (CRISPR effect) and S3 Data (Frequency of escape).*

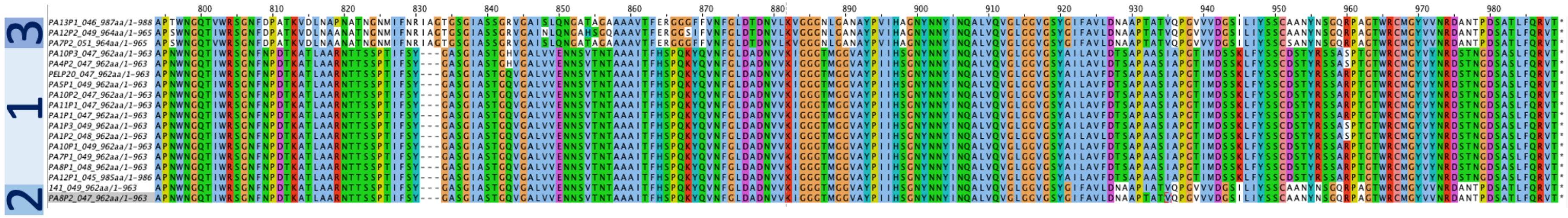

***Figure S4. Alignment of protein coding sequence for homologues of PA1P3 gp49 reveals 3 distinct sequence types.*** *The data underlying this figure can be found in S5 Data.*

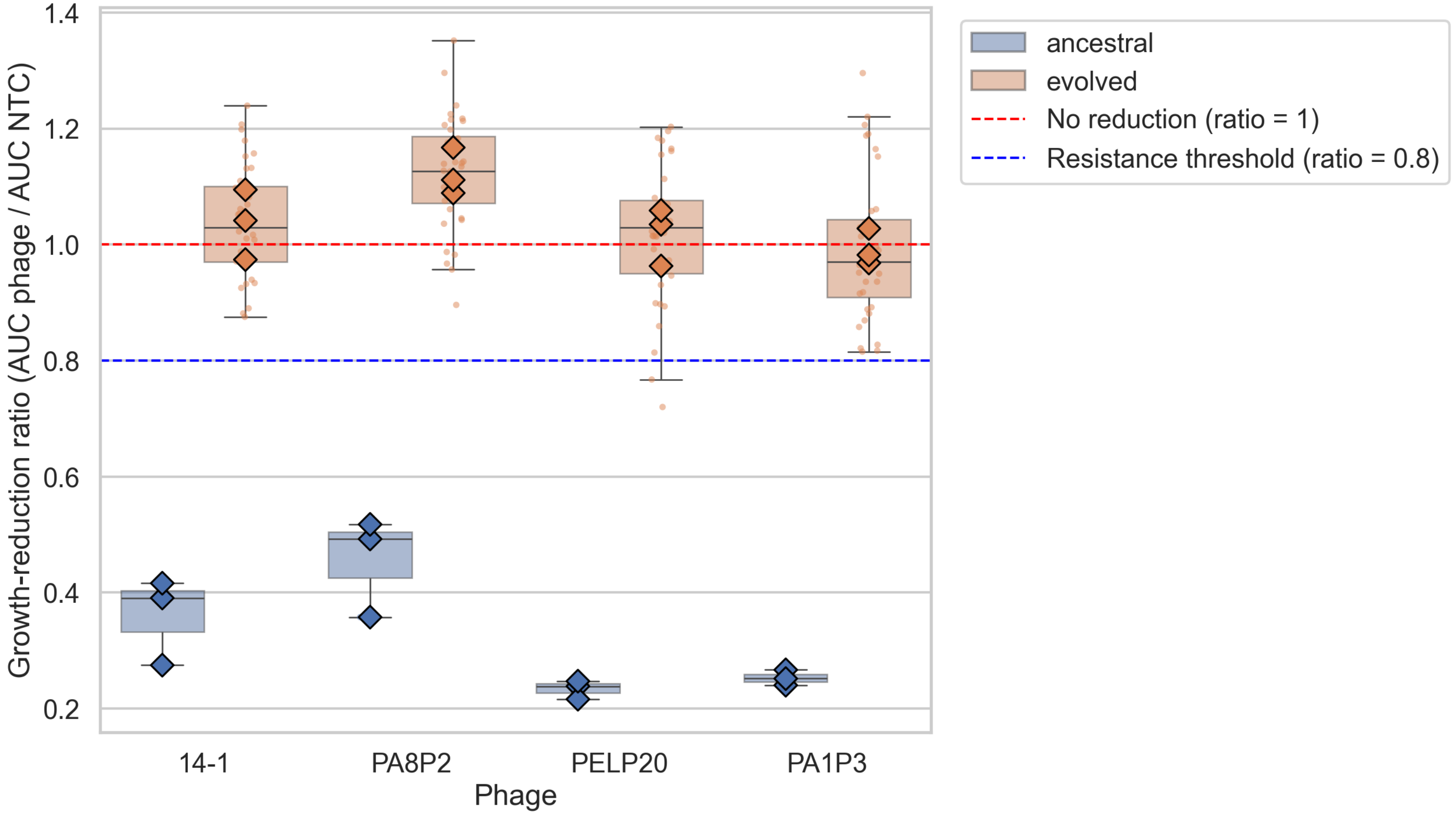


***Figure S5. Growth reduction in phage–treated ancestral and phage-evolved PA14ΔCRISPR.*** *Growth-reduction ratio (AUC (phage treated) / AUC (NTC)) for four phages (14-1, PA8P2, PELP20, PA1P3) tested against ancestral (blue) and phage-evolved (orange) bacterial populations. A ratio of 1 (red dashed line) indicates no reduction in growth relative to the untreated control; lower values indicate stronger phage-mediated suppression. Boxes show the interquartile range of individual colony-level ratios (small points); diamonds show the mean ratio for each independent biological replicate (lineage; n = 3 lineages per phage per treatment). Ancestral populations showed reduced growth relative to NTC for all four phages, while the growth reduction ratio remained above 0.8 for all but two phage-evolved clones of PELP20, indicating phage resistance. The data underlying this figure can be found in S7 Data.*

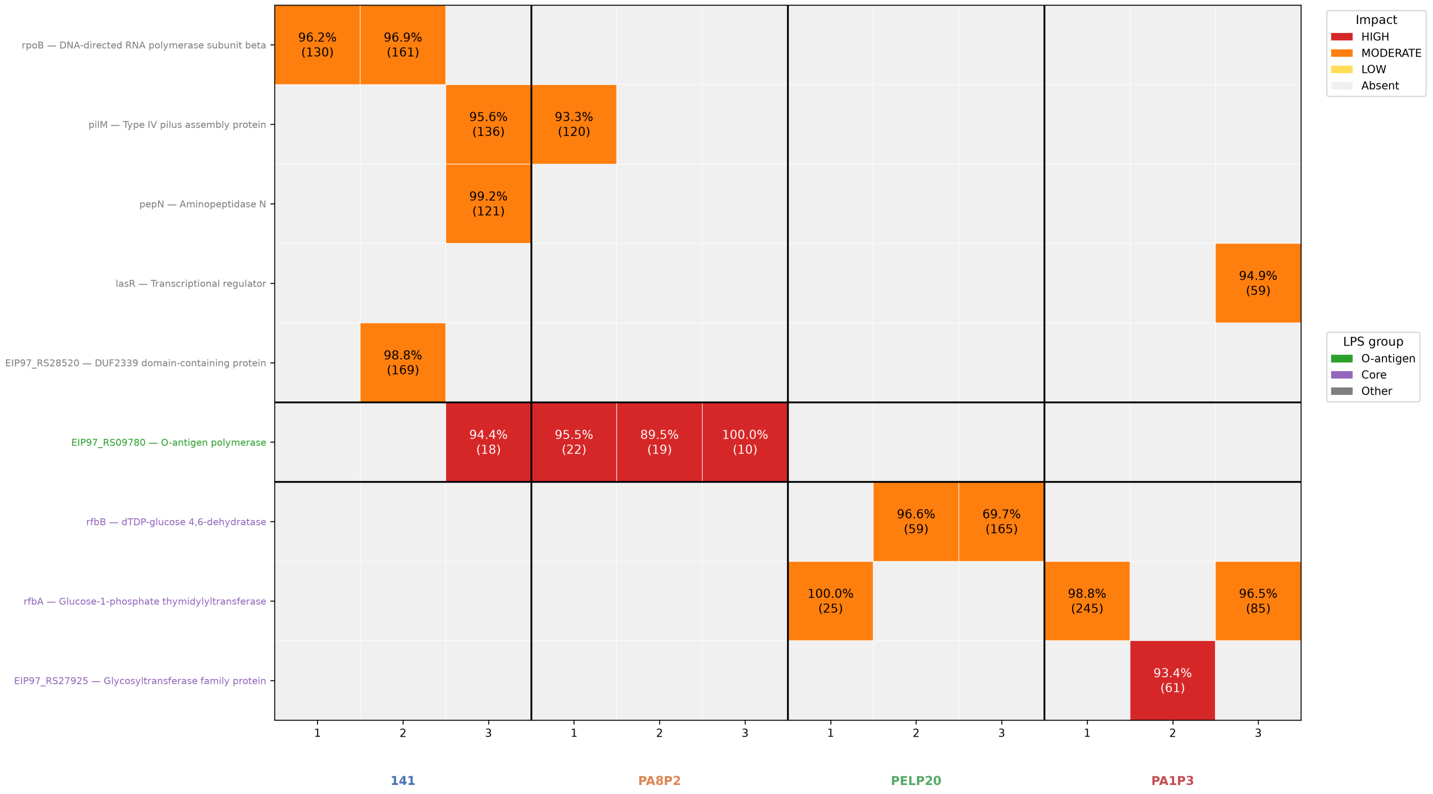


***Figure S6. Mutations detected in phage-evolved* Pseudomonas aeruginosa *lineages, by gene and predicted impact.*** *Heatmap of mutations identified (relative to the ancestral genome) in three independently evolved lineages (columns, numbered 1–3) for each of four phages used for evolution (14-1, PA8P2, PELP20, PA1P3). Rows show gene mutation presence, labeled by locus tag/gene name and annotated product. Cell colour indicates the predicted impact of the mutation in that gene (SnpEff classification): HIGH (dark red) — mutations predicted to severely disrupt protein function (e.g. frameshift, premature stop, splice-site loss); MODERATE (red-orange) — missense mutations altering the encoded amino acid; LOW (yellow-orange) — mutations with minimal predicted effect (e.g. synonymous changes); light grey — no mutation detected. Cells annotated with alternate read percentage and total coverage. Row labels coloured by predicted impact on LPS — green: O-antigen biosynthesis, purple: core oligosaccharide biosynthesis, grey: non-LPS associated loci. The data underlying this figure can be found in Table S4.*

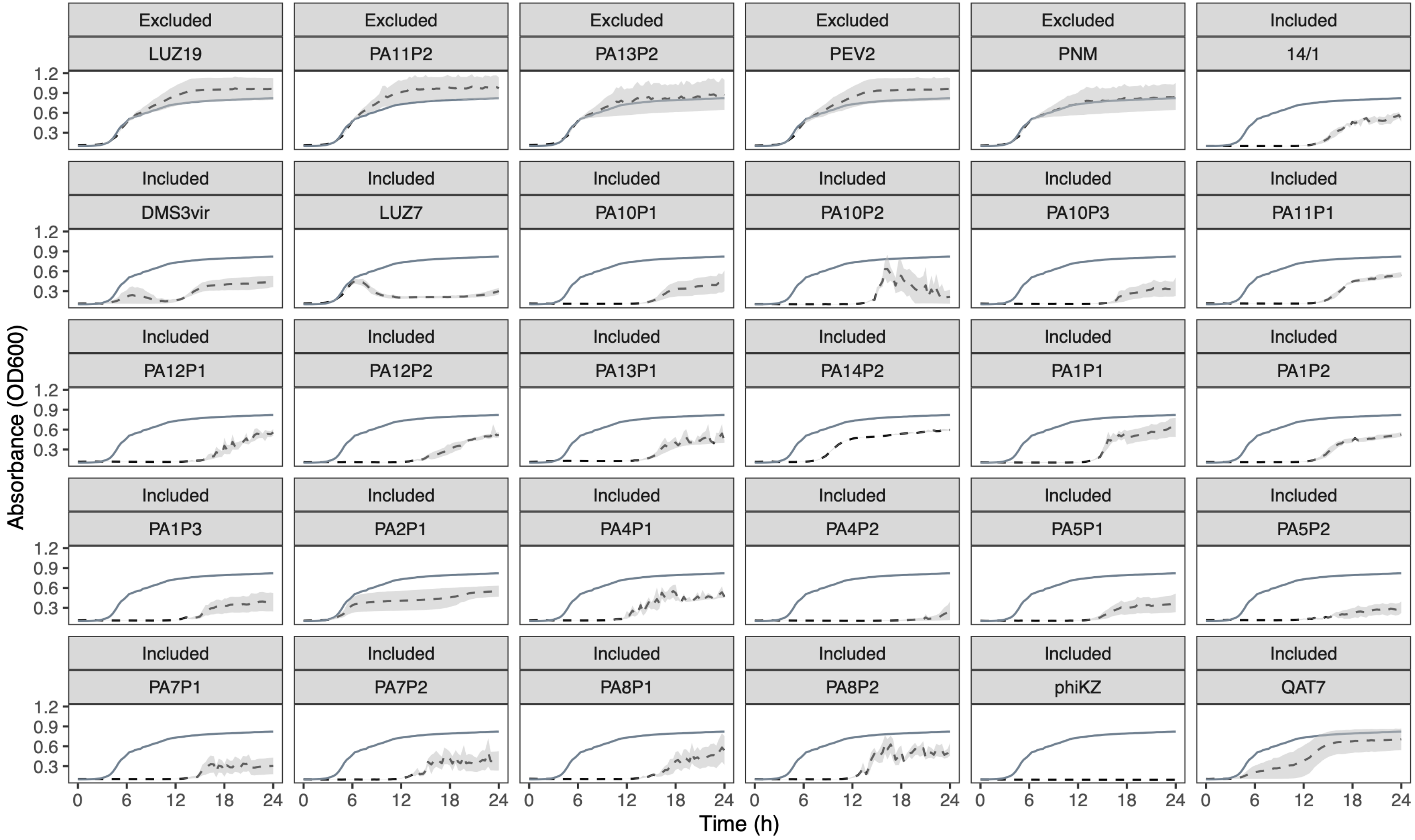


***Figure S7. Phage screening to determine infectivity in PA14 strain.*** *Growth curves show bacterial growth (PA14 CRISPR KO) in the presence (dashed line, mean of 3 replicates, shaded area shows the S.E.M.) and absence (solid line) of each phage. The data underlying this figure can be found in S8 Data.*

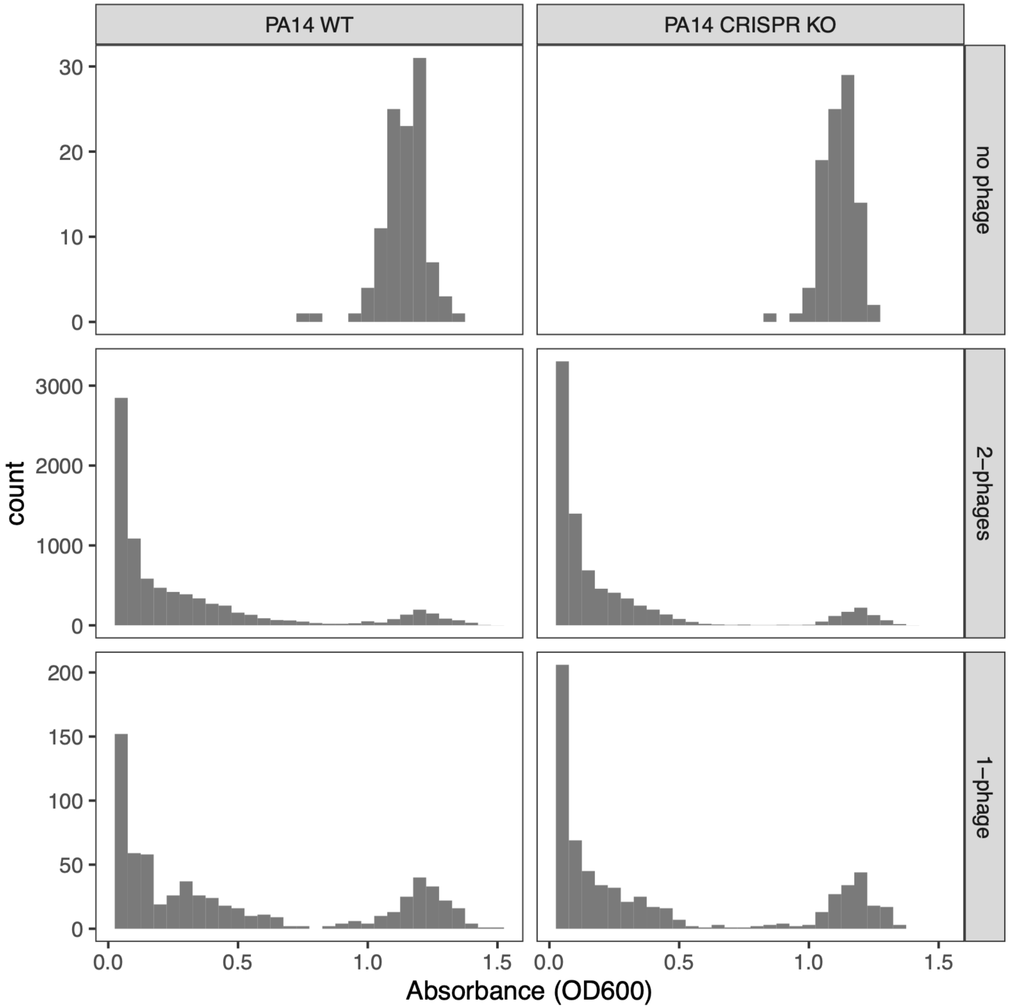


***Figure S8. Distribution of raw absorbance data used to calculate efficacy.*** *Absorbance (OD at 600nm) after 18h growth in synthetic cystic fibrosis medium against different phage treatments (rows: none, single, pairs) for PA14 either with (left panels) and without (right panels) functional CRISPR-Cas. The data underlying this figure can be found in S9 Data.*
